# Time-Resolved Mapping in Calves Reveals BoHV-1 Shift From Mucosal Replication to Trigeminal Neuroinvasion and IFN-α–bPML Antagonism

**DOI:** 10.1101/2025.08.22.671710

**Authors:** Yong Wang, Jing Cheng, Fanyu Wang, Mengyao Cao, Linyi Zhou, Wenxiao Liu, Yongqing Li

## Abstract

Bovine herpesvirus 1 (BoHV-1) causes huge cattle losses, yet the transition from mucosal replication to neuroinvasion is poorly resolved. Using a controlled calf model, we integrated quantitative virology and transcriptomics to map pathogenesis and define the role of promyelocytic leukemia protein (PML). Calves inoculated intranasally/ocularly (1.4×10^6 pfu/head) were sampled daily (1–14 dpi) for gB-qPCR; tissues at 4 and 14 dpi were analyzed for viral DNA and trigeminal ganglia (TG) mRNA-seq. Shedding peaked at 3–6 dpi (nasal > ocular ≫ rectal) and declined by 10–14 dpi. Tonsils bore the highest burden at 4 dpi; TG was low-positive at 4 dpi but remained positive at 14 dpi, indicating neuroinvasion. TG programs shifted from early proteostasis priming (4 dpi) to immune/ECM activation with synaptic repression (14 dpi). In MDBK/Vero cells, IFN-α increased bovine PML (bPML) and enlarged PML nuclear bodies (PML-NBs), reducing very-early viral DNA, whereas BoHV-1 disrupted PML-NB integrity. bPML isoforms diverged (bPML1/6 pro-viral; bPML3/4/5 antiviral). Cross-species STRING mapped a conserved PML–SUMO1–UBE2I–DAXX–SP100 core. These data delineate the mucosal-to-neuronal trajectory, establish PML as both effector and viral target, and nominate SUMO/ubiquitin-linked proteostasis as a tractable antiviral leverage point.

## 1. Introduction

BoHV-1, an alphaherpesvirus and the etiologic agent of infectious bovine rhinotracheitis, causes respiratory, ocular, and reproductive disease with substantial worldwide economic impact [1, 2]. After primary replication in the upper-respiratory mucosa, BoHV-1 establishes lifelong latency predominantly in TG sensory neurons and can reactivate under stress, driving renewed shedding and transmission [3, 4]. Transmission tightly correlates with mucosal shedding—most prominently from the nasal cavity, with ocular shedding also observed in natural and experimental settings [1]. Despite extensive descriptions of clinical disease and vaccine effects on shedding, high-resolution, quantitative characterizations that couple early shedding kinetics with parallel tissue distribution in experimentally infected calves remain limited [5, 6].

In addition to innate and adaptive immunity, host control of alphaherpesviruses depends on intrinsic nuclear defenses. A central component is PML, which assembles PML-NBs—phase-separated condensates that scaffold antiviral factors. PML-NB integrity requires SUMOylation of PML and other constituents and is enhanced by type I interferons (IFN-α/β), positioning PML at the interface of interferon signaling, proteostasis, and chromatin regulation [5, 7]. Alphaherpesviruses have evolved countermeasures: in herpes simplex virus 1 (HSV-1), the immediate-early ubiquitin ligase ICP0 localizes to PML-NBs, promotes degradation of PML and SP100, and dismantles NB architecture to relieve repression of viral gene expression [8, 9]. BoHV-1 encodes a related RING-finger protein, bICP0, which is essential for efficient productive infection and exhibits ICP0-like activities, including disruption of PML-NBs and suppression of IFN-dependent transcription, thereby antagonizing intrinsic and innate antiviral programs [10, 11].

PML exists as multiple isoforms with distinct C-terminal regions that confer isoform-specific antiviral functions and nuclear-body architectures; however, this specificity has been explored mainly in human cells and HSV models. Whether bPML isoforms differentially shape BoHV-1 replication, and how PML-linked proteostasis (SUMO/ubiquitin) and transcriptional networks are reorganized in bovine TG during early infection, remain open questions [12, 13]. Additional layers of pathogenesis include abundant expression of the latency-related (LR) RNA in latently infected sensory neurons, suggesting LR gene products help govern the latency–reactivation cycle [14, 15], and the observation that bICP0-mediated inhibition of IFN-dependent transcription is critical for virulence—BoHV-1 replicates in mice only when type I IFN signaling is ablated [16, 17].

Here, we combine an in vivo calf model with quantitative virology and systems-level host profiling to address these gaps. We define BoHV-1 shedding kinetics and tissue distribution, generate time-stamped TG transcriptomes at 4 and 14 days post-infection, and dissect the IFN-α–bPML axis together with bPML isoform-specific effects on early viral genome accumulation. We further test whether BoHV-1 antagonizes bPML by disrupting PML-NBs and map PML-centered interaction networks across species to identify conserved and bovine-specific regulators. Together, these approaches delineate a progression from robust mucosal replication to TG neuroinvasion, reveal broad immune and extracellular-matrix remodeling in TG, and position bPML as both target and effector in the host–virus interplay.

## 2. Materials and Methods

### 2.1. Ethics Statement

All animal procedures complied with the Chinese Regulations for the Administration of Laboratory Animals and the Laboratory Animal Requirements of Environment and Housing Facilities (GB14925-2012), and were approved by the Institutional Animal Care and Use Committee of the corresponding institute. Animals were monitored at least twice daily; pre-specified humane endpoints and euthanasia (pentobarbital sodium, i.v.) were applied as required. Personnel performing procedures were trained and certified in large-animal handling and biosafety.

### 2.2. Viruses, Cells, Experimental Animals, and Overall Design

#### Cells

Human embryonic kidney 293T cells and Madin–Darby bovine kidney (MDBK, ATCC CCL-22) cells were cultured in DMEM (Gibco/Invitrogen) supplemented with 10% fetal bovine serum (Hyclone), 1% L-glutamine, and 1% penicillin–streptomycin at 37 °C in 5% CO2. Cells were passaged with 0.25% trypsin–EDTA and were routinely confirmed mycoplasma-free.

#### Virus

BoHV-1 (laboratory stock) was propagated in MDBK cells (ATCC CCL-22) maintained in DMEM supplemented with 10% heat-inactivated fetal bovine serum (FBS), 1% penicillin–streptomycin, at 37 °C, 5% CO₂. Vero cells (ATCC CCL-81) were used for bPML isoform overexpression assays.

#### Animals and study design

Clinically healthy neonatal calves were acclimatized for ≥7 days and randomly assigned to control or infection groups. Calves in the infection group were inoculated intranasally and ocularly with 1.4 × 10⁶ pfu/head BoHV-1 in sterile PBS. Ocular, nasal, and rectal swabs were collected daily from 1–14 dpi for qPCR-based shedding analysis; subsets were euthanized at 4 dpi and 14 dpi for tissue collection (e.g., palatine tonsil, trigeminal ganglion, upper-airway/lymphoid tissues) and downstream virologic and transcriptomic analyses.

### 2.3. Virus Propagation and Titration

BoHV-1 working stocks were generated by infecting near-confluent MDBK monolayers at multiplicity of infection (MOI) = 1 in serum-free DMEM for 1.5 h with intermittent rocking. After adsorption, cells were washed with PBS and incubated in maintenance medium (2% FBS) until cytopathic effect (CPE) was extensive. Supernatant and cells were freeze–thawed three times, clarified (3,000 × g, 10 min), aliquoted, and stored at −80 °C. Infectious titer was determined on MDBK cells in 96-well plates by endpoint dilution, and TCID₅₀ values calculated by the Kärber method.

### 2.4. Calf Inoculation, Swab Sampling, and Tissue Processing

Calves were inoculated as above; swabs (ocular, nasal, rectal) were collected using sterile polyester swabs and placed into 1 mL PBS (on ice), vortexed, and stored at −80 °C until extraction. At necropsy (4 and 14 dpi), tissues (tonsil, TG, additional URT/lymphoid organs) were dissected aseptically, partitioned for DNA/RNA, snap-frozen in liquid nitrogen, and stored at −80 °C.

### 2.5. DNA Extraction and Absolute qPCR for BoHV-1 gB

Swabs. Tubes were vortexed (30 s), incubated at room temperature (1 h), and centrifuged (12,000 rpm, 8 min). 200 µL supernatant was used for DNA extraction (DNeasy Blood & Tissue Kit, Qiagen) and eluted in 40 µL nuclease-free water.

Tissues. ∼20–30 mg tissue was homogenized in ATL buffer with stainless-steel beads (tissue lyser), followed by DNA extraction (DNeasy).

Absolute qPCR. Viral DNA was quantified by absolute qPCR targeting gB using plasmid standards spanning 10⁸–10¹ copies per reaction to construct a standard curve (CFX96, Bio-Rad; 20 µL reaction with SYBR Green master mix; 95 °C 2 min; 40 cycles of 95 °C 15 s, 58–60 °C 60 s; melt-curve check). Data are expressed as gB copies per swab or normalized to input tissue DNA (ng). Primer sequences and efficiencies are listed in Supplementary Table S1.

### 2.6. RNA Isolation, Library Construction, and RNA-seq

TG samples (control, 4 dpi, 14 dpi; n = 3 per group) were homogenized in TRIzol (Invitrogen). Poly(A)⁺ mRNA was enriched with oligo(dT) magnetic beads, reverse-transcribed, and converted to double-stranded cDNA; libraries were prepared according to the manufacturer’s instructions and sequenced on an Illumina NovaSeq 6000 (paired-end). Raw reads were quality-filtered with fastp and aligned to the bovine reference genome using HISAT2 and TopHat2; transcripts were quantified to TPM for exploratory QC.

### 2.7. Differential Expression and Functional Enrichment

Differentially expressed genes (DEGs) were identified using DESeq2 with default normalization and Wald tests. Unless otherwise specified, thresholds were FDR (Benjamini–Hochberg) < 0.05 and |log₂FC| ≥ 1. GO (Biological Process/Cellular Component/Molecular Function) and KEGG over-representation analyses were performed on DEG sets; dot plots display gene counts and adjusted *P*. EggNOG/COG functional categories were assigned to DEGs to assess category-level dynamics across 4 dpi and 14 dpi.

### 2.8. Gene Set Enrichment Analysis (GSEA)

A pre-ranked GSEA was performed (MSigDB/GO collections) by ranking all genes using a signed statistic derived from differential expression (e.g., log₂FC × −log₁₀*P*). Enrichment was summarized by normalized enrichment score (NES) and FDR q-value. Leading-edge genes were visualized by Z-scored heatmaps. Analysis parameters and permutation settings are detailed in the figure legends and Methods.

### 2.9. Generation of bPML-Modified Cell Lines and Isoform Expression

#### 2.9.1. Plasmids and Reagents

The lentiviral expression vector pTK643.EF1α-IRES-GFP and packaging plasmids Δ NRF and VSV-G were gifts from Prof. Liguo Zhang. The knockdown backbone pLKO.1-puro was obtained from Addgene. Transfections used Lipofectamine 2000 (Invitrogen) according to the manufacturer’s instructions.

#### 2.9.2. shRNA Design and Cloning

Short hairpin RNAs (shRNAs) targeting bovine PML were designed using Thermo Fisher’s siRNA design resource and synthesized by Tsingke (Beijing). Annealed, double-stranded oligonucleotides were cloned into pLKO.1-puro using AgeI/EcoRI restriction sites and T4 DNA ligase. Oligo sequences and target coordinates are provided in Supplementary Table S2.

#### 2.9.3. Lentivirus Production and Transduction

Third-generation lentiviruses were produced by co-transfecting HEK293T cells with Δ NRF and VSV-G together with either pLKO.1-bPML shRNA (knockdown) or pTK643-EF1α-bPML-IRES-GFP (overexpression). Supernatants were collected at 48 h, clarified, and filtered (0.45 µm). For transduction, 1×10^6 MDBK cells were exposed to 1 mL lentivirus with 8 µg/mL polybrene and spinoculated at 3,000 rpm for 3 h. Inoculum was replaced with complete medium and cells were recovered for 48–72 h prior to selection or sorting.

#### 2.9.4. Generation of Stable Knockdown and Overexpression Lines

For PML knockdown, MDBK cells were selected in 4 µg/mL puromycin to establish shPML-MDBK lines; a non-targeting shRNA served as control. For PML overexpression, MDBK (or 293T) cells transduced with pTK643-EF1α-bPML-IRES-GFP were enriched by FACS (BD Aria III) for GFP-positive cells to derive PMLHi-MDBK (or 293T-bPML) lines. All lines were validated by qRT-PCR, immunoblotting, and immunofluorescence for bPML expression and PML-NB features.

Knockdown/overexpression in MDBK. Stable shPML-MDBK (bPML knockdown) and PML^Hi-MDBK (bPML overexpression) derivatives were generated by lentiviral transduction (standard third-generation packaging), followed by antibiotic selection and validation by qRT-PCR and immunoblotting. Immunofluorescence (IF) was used to assess PML nuclear-body (PML-NB) abundance/organization.

Isoform overexpression in Vero. Plasmids encoding bPML1–bPML6 (C-terminal isoform diversity) or empty vector were transfected into Vero cells (lipid-based reagent). Expression was verified by immunoblot; cells were used 24 h post-transfection for infection assays.

### 2.10. Recombinant bPML expression and purification (prokaryotic system)

Bovine PML cDNA fragments (e.g., 1174–1279 bp, 1365–1800 bp; GenBank OM324027) were synthesized (Tsingke) and cloned into pET-32a(+). Constructs were transformed into Transetta (DE3). Cultures were induced with 0.5–1 mM IPTG at 25 ° C for ∼7 h. Cells were lysed in 20 mM Tris–HCl pH 8.0, 150 mM NaCl; clarified lysates were applied to Ni-NTA resin (Thermo Fisher) and eluted per manufacturer’s instructions. Fractions were analyzed by SDS–PAGE and Coomassie staining; purity was assessed prior to downstream immunization or validation.

### 2.11. Antibody production and validation

New Zealand white rabbits and BALB/c mice were immunized with purified recombinant bPML according to standard schedules (prime + multiple boosts with adjuvant). For monoclonal antibody generation, splenocytes from immunized BALB/c mice were fused with SP2/0 myeloma cells, and hybridomas were screened by ELISA/IFA/Western for bPML specificity, yielding clone bPML-2G5. Validation employed transient transfection of pCMV-HA-C-bPML into 293T cells followed by IF (bPML-2G5 primary, appropriate secondary; DAPI nuclear counterstain) and immunoblot of whole-cell lysates. Antibodies detailed information are provided in Supplementary Table S3.

### 2.11. IFN-α Stimulation, Immunoblotting, and Immunofluorescence

MDBK cells were treated with recombinant IFN-α (dose and duration in figure legends), lysed in RIPA buffer with protease/phosphatase inhibitors, and probed by immunoblot with anti-bPML mAb (bPML-2G5) and loading controls. For cell IF, cells on coverslips were fixed (paraformaldehyde), permeabilized (0.1% Triton X-100), blocked, and incubated with anti-bPML and appropriate secondary antibodies; nuclei were counterstained with DAPI. For tissue IF, TG sections (4 and 14 dpi) underwent antigen retrieval, blocking, and double IF with anti-bPML and anti-ICP0, followed by confocal imaging under matched acquisition settings.

### 2.12. Early Infection Kinetics Assay (gB qPCR)

MDBK derivatives (shPML, parental, PML^Hi) or Vero cells expressing individual bPML isoforms were infected with BoHV-1 (MOI = 1). At 0.5, 1, 2, 4, 6, and 12 hpi, total DNA was extracted and viral gB copies were quantified by qPCR to assess very-early genome accumulation. Results were normalized to cell input and analyzed across time points.

### 2.13. Disruption of PML-NBs by BoHV-1

To interrogate viral counteraction of PML restriction, MDBK cells were infected (MOI = 1) and fixed at CPE onset for double IF (bPML/ICP0). In parallel, TG sections from 4 and 14 dpi calves were analyzed by double IF to visualize PML-NB integrity and subcellular distribution. Image acquisition parameters and antibody details are provided with the figures.

### 2.14. Protein–Protein Docking

The bPML–bICP0 interface was modeled using HADDOCK with default restraints and explicit solvent refinement. Clusters were ranked by HADDOCK score; interfacial properties (e.g., buried surface area) were computed for the top cluster. Parameterization and evaluation criteria are given in the figure legend and Methods.

### 2.15. STRING Protein–Protein Interaction (PPI) Networks

STRING (v11.x) was used to build PML-centered first-shell networks for Homo sapiens and Bos taurus under high-confidence settings (evidence from curated databases, experiments, co-expression, and text mining). The canonical PML–SUMO1–UBE2I–DAXX–SP100 core and species-selective nodes were examined; combined scores determined edge thickness. Enrichment analyses followed STRING default multiple-testing correction.

### 2.16. Quantitative RT-PCR

Total RNA was extracted with TRIzol; 1 µg RNA was reverse-transcribed (random hexamers/oligo-dT mix). qRT-PCR used SYBR chemistry (CFX96) with gene-specific primers (Supplementary Table S1). Expression was normalized to GAPDH using the 2⁻ΔΔCt method. Primer specificity was confirmed by melt-curve analysis.

### 2.17. Statistics and Reproducibility

Unless specified otherwise, data are mean ± SD from ≥3 independent experiments/animals. Two-way ANOVA with post hoc multiple-comparison correction was used for time-course assays; unpaired t-tests (or Mann–Whitney) for two-group comparisons. RNA-seq multiple testing used Benjamini–Hochberg FDR. Significance threshold was P < 0.05 (two-sided). Analyses were performed in GraphPad Prism and R (v4.x) with defined seeds for stochastic procedures. Sample sizes, exact *n*, and tests are indicated in figure legends.

## 3. Results

### 3.1. Replication dynamics and early tissue distribution of BoHV-1 in calves

Early mucosal replication was assessed by serial swabbing. At 2 dpi (Supplementary Figure 1A–B), viral genomic DNA was detectable in ocular and rectal swabs. By 3 dpi (Supplementary Figure 1C), nasal swabs exhibited a sharp increase in viral copies, indicating robust replication and shedding through the upper respiratory tract. To define early tissue distribution, we quantified viral DNA (gB qPCR) in tissues collected at 4 and 14 dpi (Figure 1A–F). At 4 dpi, BoHV-1 DNA was readily detectable in upper-respiratory/lymphoid tissues, with the highest copy numbers in tonsils; lower amounts were detected in TG and other tissues (Figure 1G). By 14 dpi, viral DNA declined in tonsils relative to 4 dpi, whereas TG remained consistently positive, indicating neuroinvasion and maintenance of viral DNA in sensory ganglia during the late acute phase (Figure 1H). Additional mucosal and lymphoid tissues were positive at both time points, with greater between-tissue heterogeneity at 14 dpi. Overall, these data support a sequence of robust replication at mucosal/lymphoid sites followed by transport to TG and early establishment of persistence.

**Figure 1.**
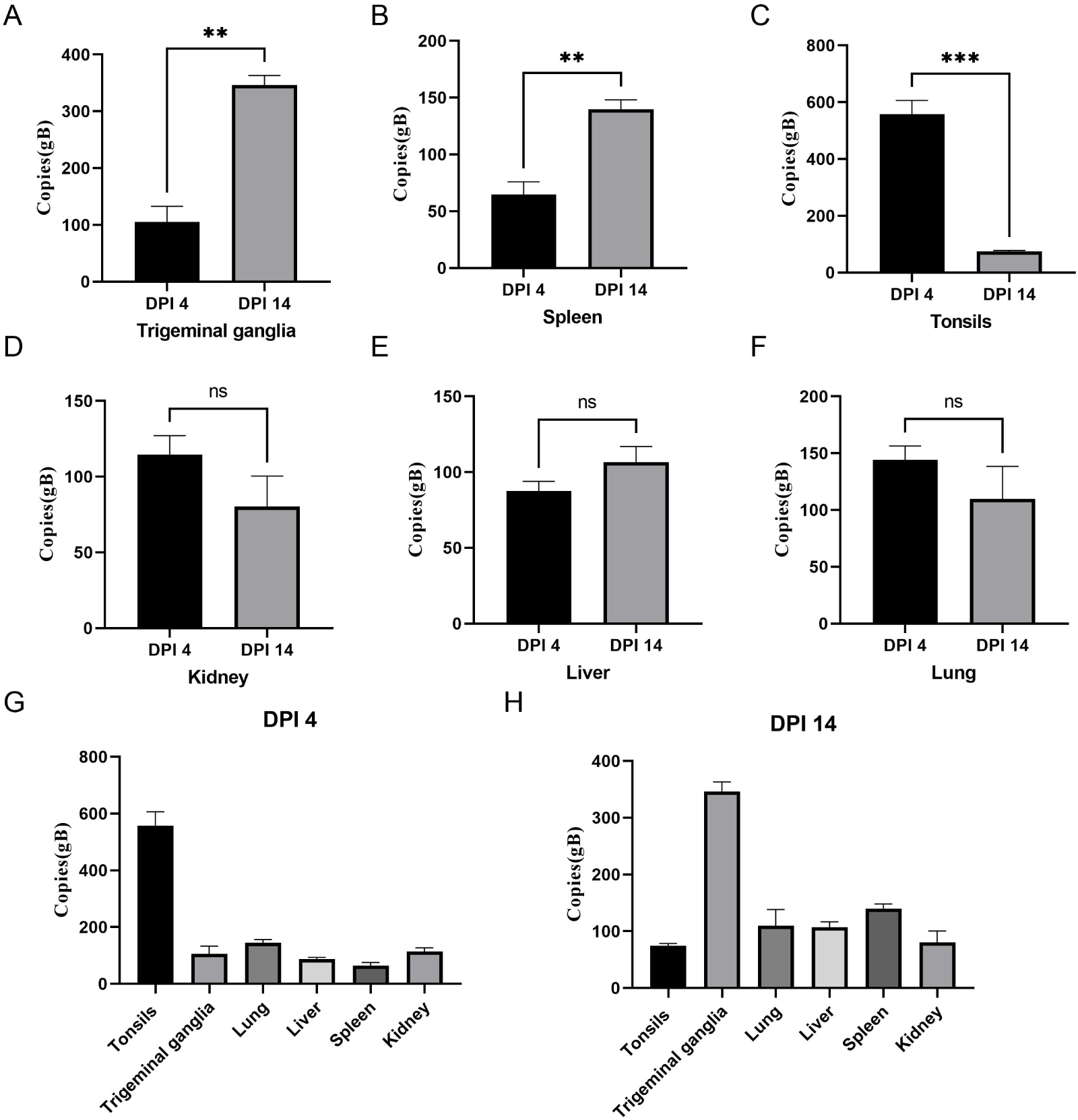
Early tissue distribution of BoHV-1 DNA in experimentally infected calves. Calves were inoculated intranasally and ocularly with BoHV-1 (1.4 × 10^6 pfu per head) and euthanized at 4 or 14 days post-infection (dpi). Palatine tonsil (A), trigeminal ganglion (TG) (C), and additional upper-airway/lymphoid tissues (B, D–F) were collected, total DNA extracted, and viral genomes quantified by absolute qPCR targeting gB. Bars show gB copy numbers normalized to input tissue DNA for each animal; horizontal lines denote group mean ± SD (n = 3 calves per time point). Statistical methods and significance criteria are detailed in Methods.

### 3.2. Time-stamped TG transcriptomes reveal early sensing signature and late immune/ECM remodeling during BoHV-1 infection

We performed mRNA-seq on TG from control calves (CG) and infected calves at 4 and 14 dpi (DPI4, DPI14; n = 3/group). Total RNA was extracted with TRIzol, poly(A)^+ RNA was enriched with oligo-dT magnetic beads, converted to double-stranded cDNA, and libraries were sequenced on an Illumina NovaSeq 6000. Reads were quality-filtered (fastp) and aligned to the bovine reference genome (HISAT2/TopHat2). TPM-normalized expression distributions were comparable across samples without apparent batch effects (Figure 2A).

**Figure 2.**
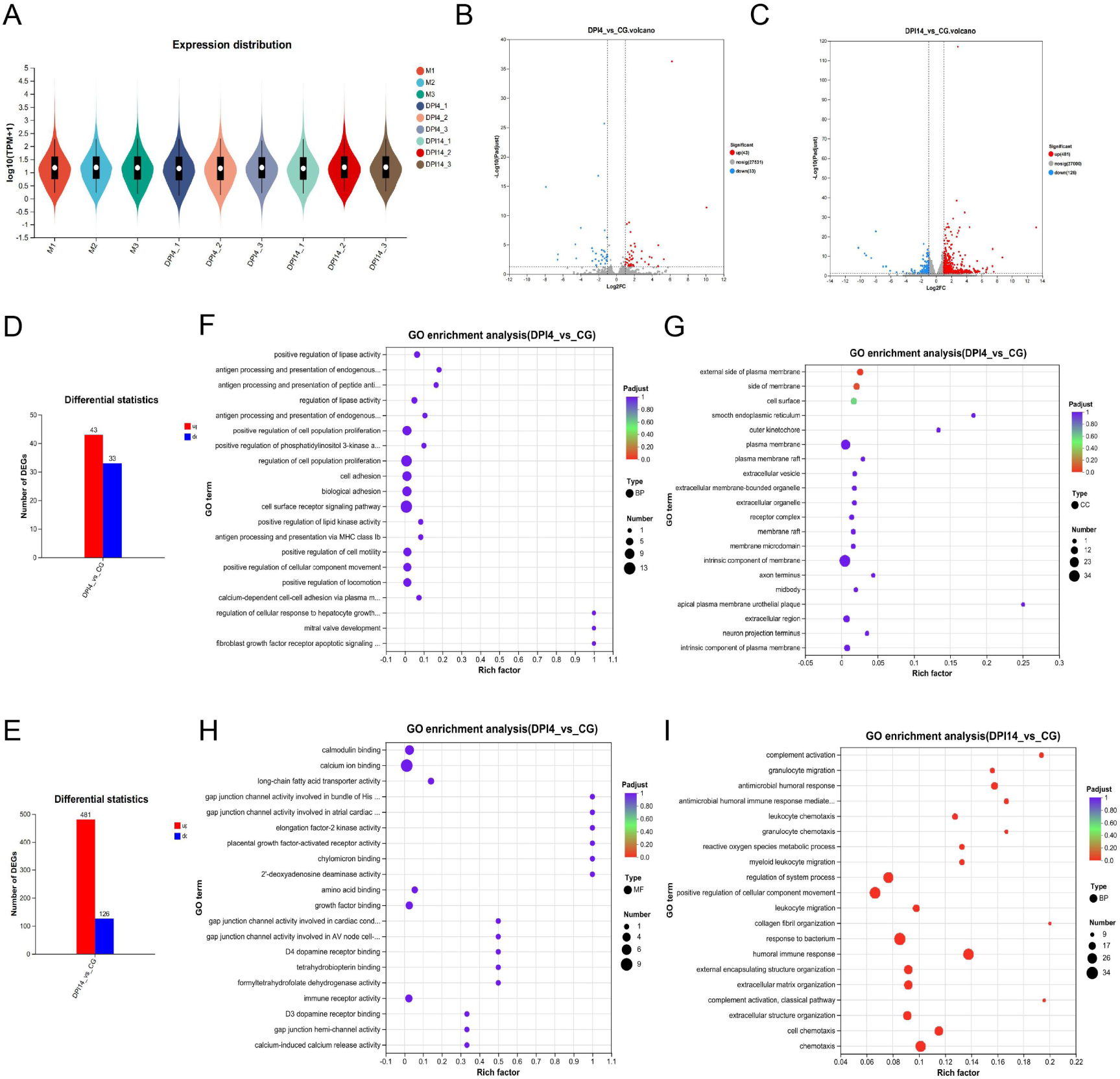
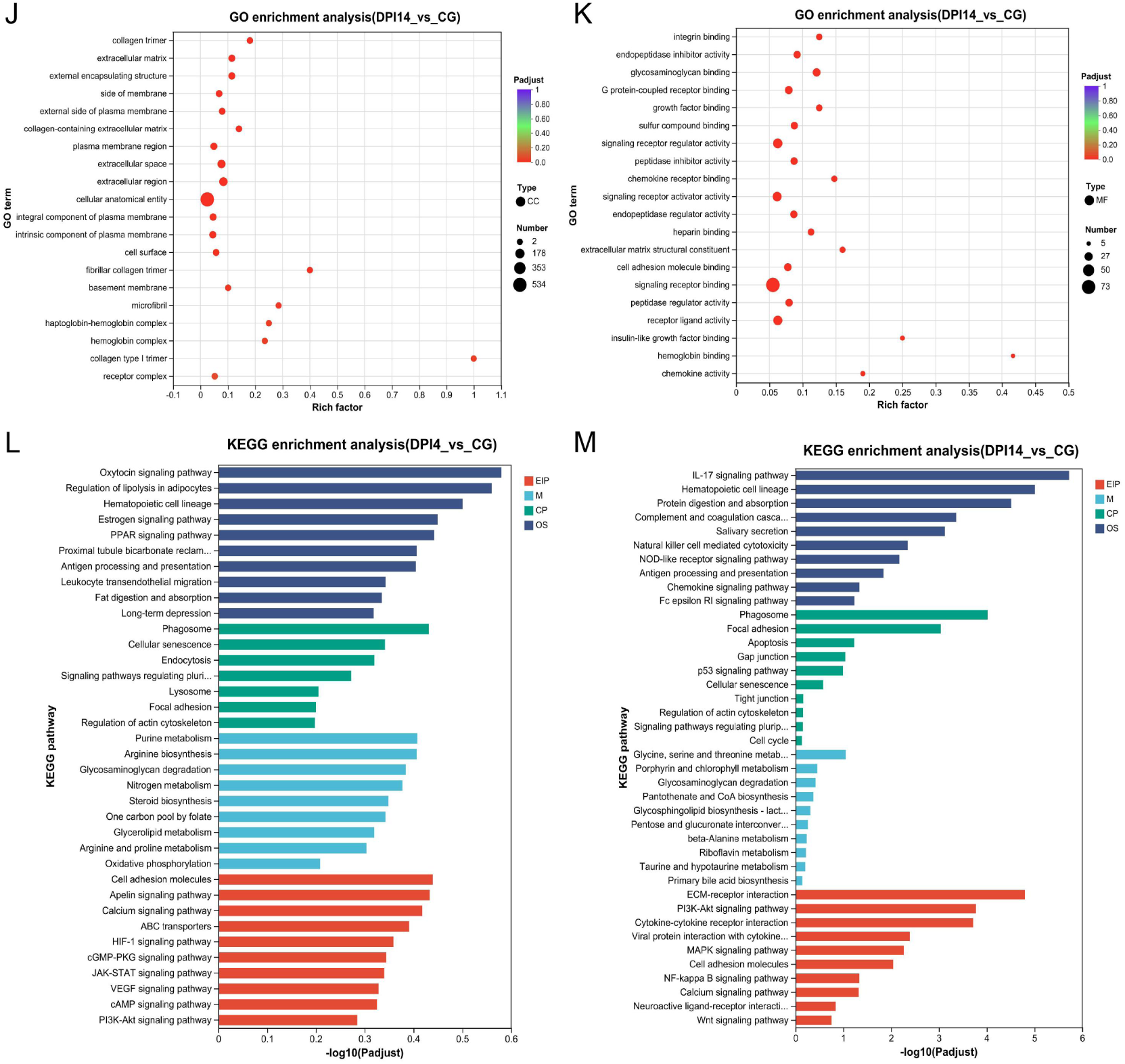
Time-resolved transcriptomic remodeling in trigeminal ganglia after BoHV-1 infection. TG from control (CG), 4 dpi (DPI4), and 14 dpi (DPI14) calves (n = 3/group) were processed for mRNA-seq; read processing, mapping, and statistics are detailed in Methods. (A) Violin plots of log10(TPM+1) indicate comparable global expression distributions across groups. (B–C) Volcano plots for DPI4 vs CG and DPI14 vs CG with log2 fold-change and adjusted-*P* thresholds indicated. (D, H) Numbers of differentially expressed genes (DEGs) at DPI4 and DPI14. (E–G) GO enrichment for DPI4 DEGs highlights an early, limited program consistent with immune sensing and cell-adhesion/motility. (I–K) GO enrichment for DPI14 DEGs shows broad activation of innate/humoral immunity with extracellular-matrix remodeling and reduced neuronal/synaptic features. (L–M) KEGG over-representation analysis: DPI4 emphasizes endocytosis/trafficking and signaling priming, whereas DPI14 is dominated by inflammatory/antiviral pathways and apoptosis/ECM modules. Dot size reflects gene counts; color encodes Benjamini–Hochberg adjusted *P*.

Relative to CG, transcriptional changes at 4 dpi were modest: 76 DEGs (43 up, 33 down) (Figure 2B,D). GO enrichment highlighted antigen processing/presentation, cell adhesion/motility, and regulation of lipid-kinase–linked signaling (Figure 2F); cellular-component terms were enriched for the plasma membrane (including membrane rafts), receptor complexes, and extracellular vesicles (Figure 2G); molecular-function terms included calcium/calmodulin binding and immune-receptor activity (Figure 2H). KEGG analysis suggested early engagement of endocytosis/lysosome–phagosome pathways, leukocyte transendothelial migration, and signaling via JAK–STAT, PI3K–Akt, and second-messenger cascades (cAMP/cGMP/calcium) (Figure 2L).

By 14 dpi, TG exhibited broad transcriptional reprogramming with 607 DEGs (481 up, 126 down) (Figure 2C,E). GO biological-process terms were dominated by complement activation, granulocyte chemotaxis, antimicrobial humoral responses, leukocyte chemotaxis, and reactive-oxygen-species metabolic processes (Figure 2I); cellular-component terms were enriched for extracellular matrix (ECM), collagen-containing ECM, and basement membrane (Figure 2J); molecular-function terms included integrin binding, chemokine/cytokine receptor binding, and peptidase inhibitor/regulator activity (Figure 2K). KEGG pathways indicated robust activation of antiviral and inflammatory signaling—IL-17, complement and coagulation cascades, NK cell–mediated cytotoxicity, NOD-/RIG-I-like receptor, chemokine, Toll-like receptor, NF-κB, MAPK, and TNF pathways—together with apoptosis, p53, gap junction, and ECM–receptor interaction (Figure 2M). Collectively, TG shows early, limited priming at 4 dpi, followed by widespread innate/humoral immune activation and ECM remodeling at 14 dpi, consistent with neuroinvasion and a sustained antiviral/inflammatory milieu.

### 3.3. EggNOG/COG and GSEA reveal late-phase immune activation and synaptic downregulation in BoHV-1–infected TG

EggNOG/COG classification revealed a time-dependent pattern: at 4 dpi, enrichment in K/O/T/I categories was consistent with early sensing and proteostasis priming (Figure 3A,C), whereas at 14 dpi DEGs were dominated by V/T/O categories, together with D/L/U/Z, reflecting defense, signaling, PTM/turnover, cell cycle/repair, intracellular trafficking/secretion, and cytoskeleton (Figure 3B,D–G). GSEA comparing 14 dpi with controls showed positive enrichment of G-protein–coupled receptor binding, cell chemotaxis, antimicrobial humoral response, receptor complex, cell–cell signaling, and secretion (typical NES ∼1.55–1.74; FDR as shown; Figure 4A–G). Negatively enriched sets included nucleoside/guanyl nucleotide binding, RNA polymerase II–related transcriptional activation, and regulation of peptidase activity (e.g., NES ∼ −1.52; FDR ∼ 0.27), with down-regulation of synaptic receptor/neurotransmission modules (e.g., GABRG1, CHRNA3, GRID1, GRIK3, DLGAP1; Figure 4H). Thus, at 14 dpi TG displays immune/chemotactic activation with ECM remodeling alongside suppression of neuronal/synaptic programs.

**Figure 3.**
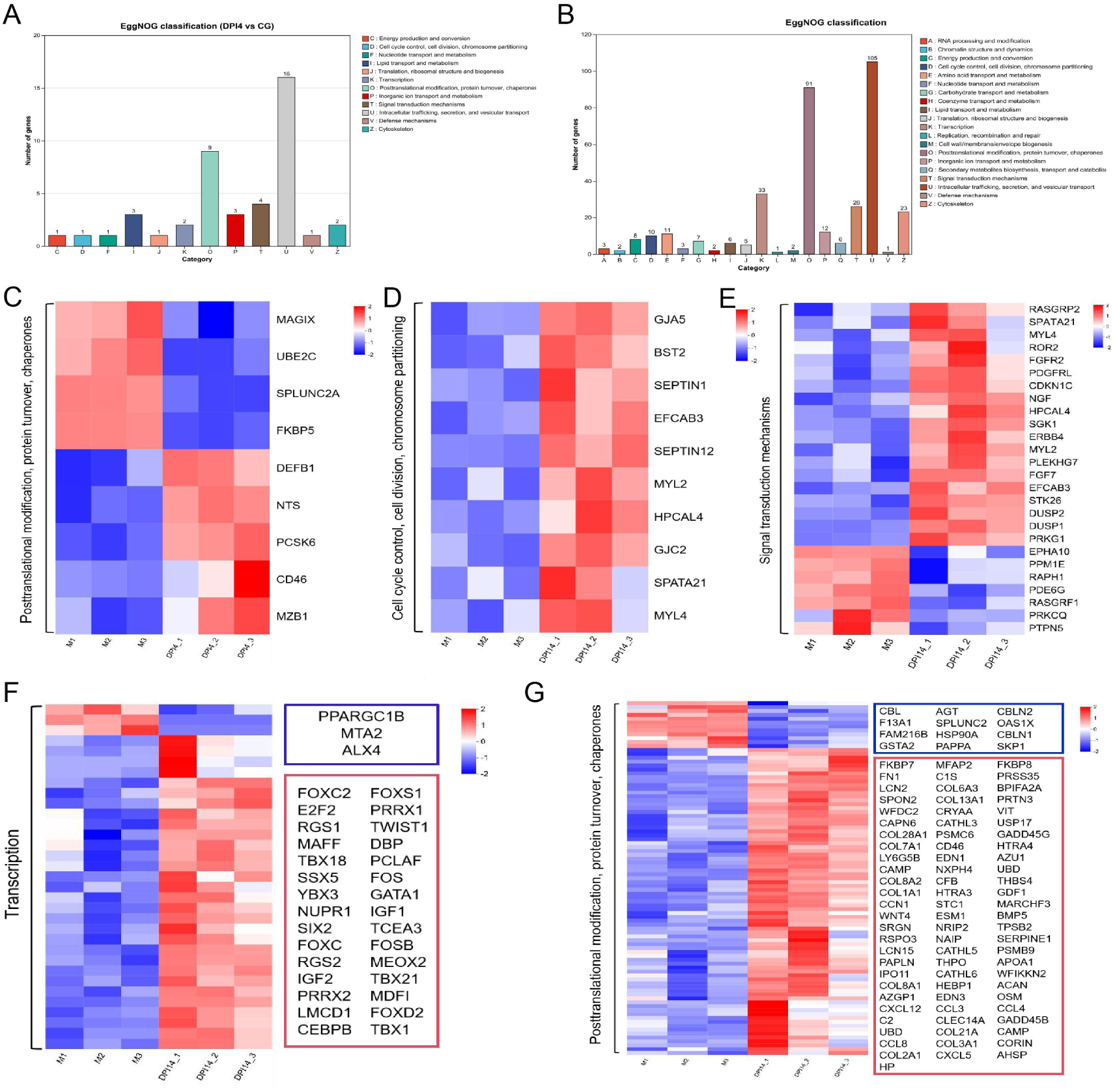
EggNOG/COG classification and representative gene modules in TG after BoHV-1 infection. TG RNA-seq from control (CG), 4 dpi (DPI4), and 14 dpi (DPI14) calves (n = 3/group) was analyzed for differential expression and functional annotation as described in Methods. (A–B) DEGs (DPI4 vs CG; DPI14 vs CG) were categorized by EggNOG/COG. (C–E) Heatmaps of selected transcription/signaling/metabolic modules showing progressive shifts from CG to DPI4 and DPI14. (F–H) Heatmaps of immune and protein-turnover modules with dominant up-regulation at DPI14. Color scale indicates Z-scored expression; statistical thresholds are detailed in Methods.

**Figure 4.**
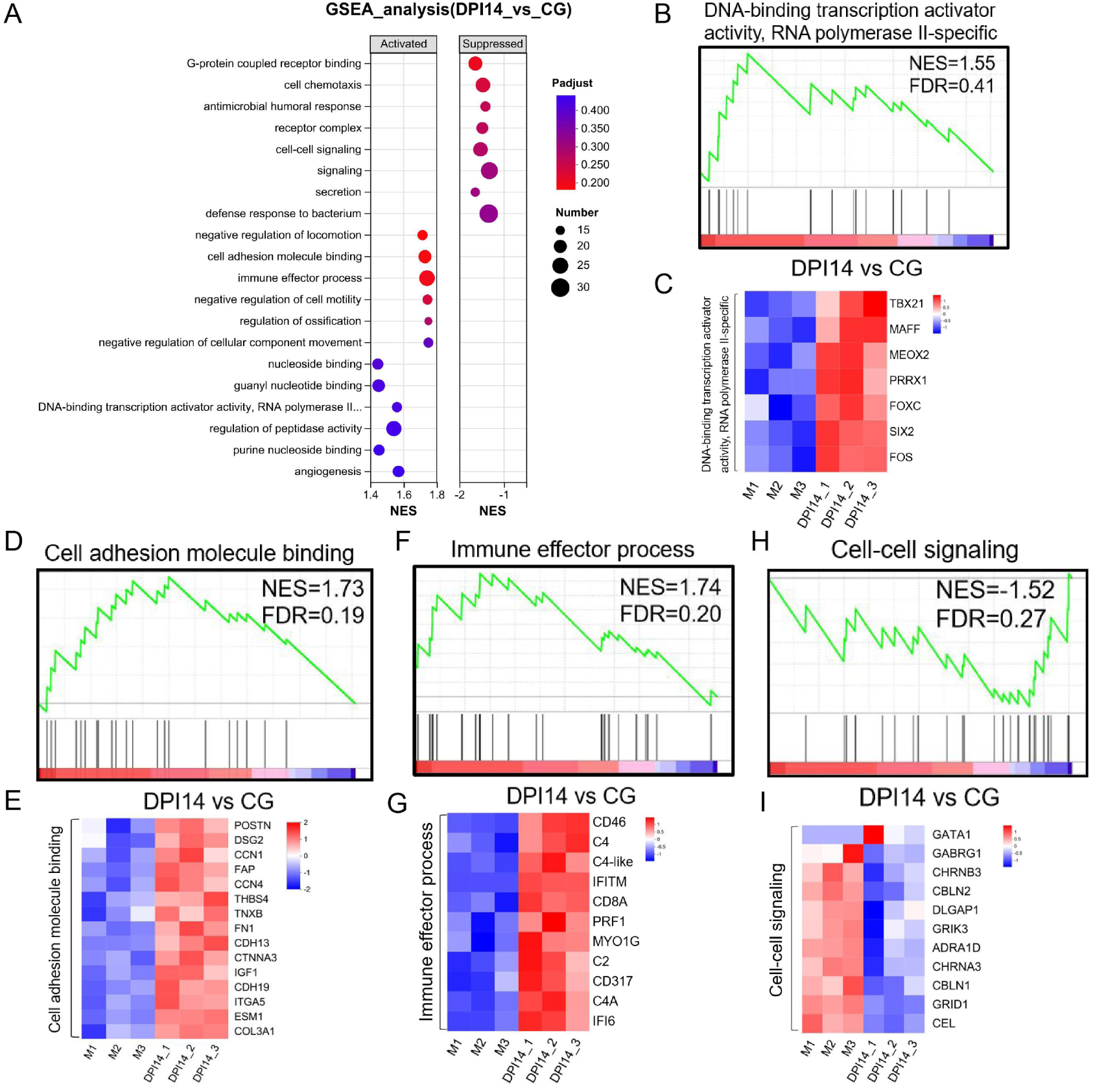
Gene set enrichment analysis (GSEA) of trigeminal ganglia at 14 dpi versus controls. RNA-seq data from TG (n = 3 per group) were analyzed by pre-ranked GSEA comparing 14 dpi to control, using a signed differential statistic to rank all genes and MSigDB/GO collections as gene sets (multiple-testing controlled as in Methods). (A) Dot plot summarizing representative activated and suppressed gene sets with normalized enrichment scores (NES), FDR-adjusted P values, and set sizes. (B–D) Enrichment plots with corresponding leading-edge heatmaps for exemplar activated programs. (E) Enrichment plot and leading-edge heatmap for a representative suppressed program. Heatmaps display Z-scored expression across samples; thresholds, databases, and permutation settings are detailed in Methods.

### 3.4. Conserved PML–SUMO nuclear-body core with species-specific extensions in human and bovine interactomes

STRING analysis revealed densely interconnected PML-centered networks in human and bovine datasets (Figure 5A,B). In each species, PML sits at the hub of a canonical PML-NB module (PML–SUMO1–UBE2I–DAXX–SP100), indicating conservation of SUMO-dependent nuclear-body architecture. Networks converged on TP53–MDM2 and included RARA, RUNX1, and RNF4, linking PML to chromatin control, cell-cycle/stress signaling, and SUMO-triggered proteasomal turnover. Species-selective differences were evident: the human network contained SPI1 (PU.1), whereas the bovine network incorporated EP300 and an unannotated protein (ENSBTAP00000065305) connected to SUMO1/UBE2I/MDM2. These features suggest a conserved core with lineage-specific extensions that may tune chromatin states and innate outputs.

**Figure 5.**
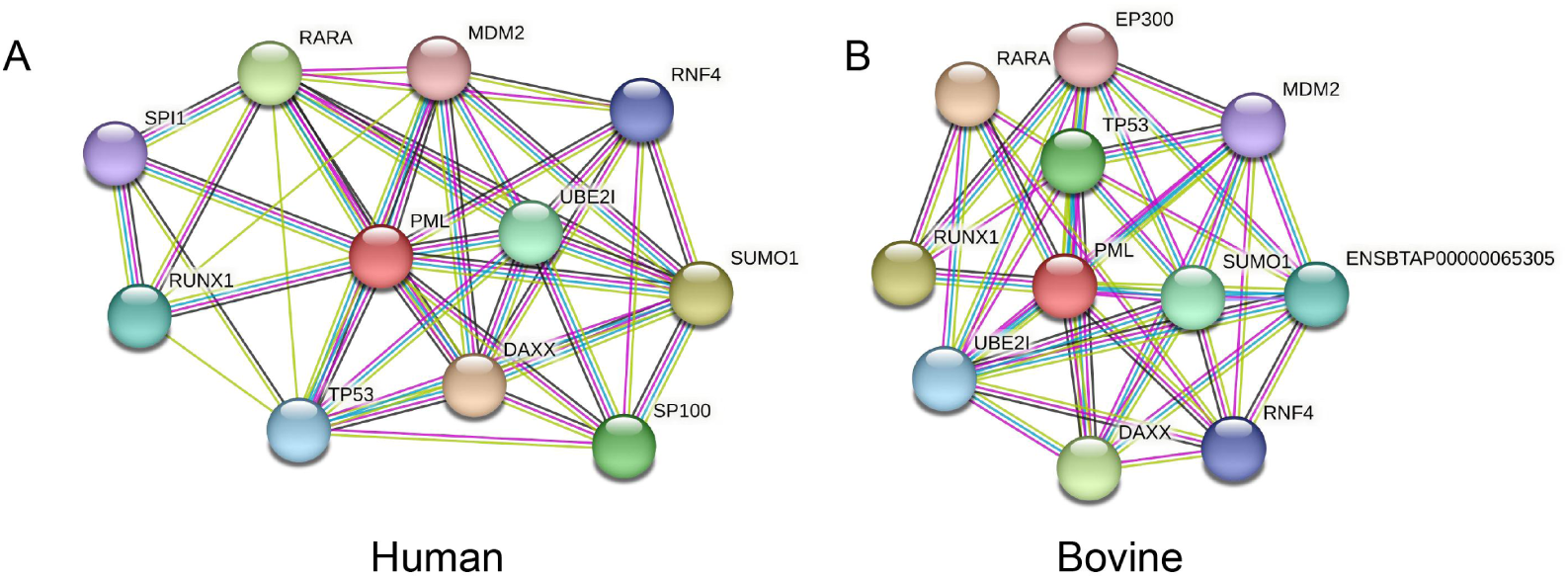
PML-centered protein–protein interaction networks in human and bovine generated with STRING. Networks were constructed in STRING using PML from Homo sapiens (A) and Bos taurus (B) as seed proteins, retrieving first-shell interactors under a high-confidence evidence model (curated databases, experiments, co-expression, and text mining). Nodes represent proteins; edges depict STRING combined-score associations (edge thickness reflects confidence). Layout, evidence channels, and enrichment procedures are described in Methods.

### 3.5. IFN-α–bPML axis suppresses very-early replication via PML-NBs

Because PML participates in IFN-α–mediated innate immunity [7], we examined whether IFN-α regulates bPML in MDBK cells. IFN-α stimulation markedly increased bPML protein (Western blot; Figure 6A), and immunofluorescence revealed pronounced remodeling of PML-NBs, with clear increases in both number and size (Figure 6B). To assess function, we generated shPML-MDBK (knockdown) and PML^Hi-MDBK (overexpression) derivatives. qPCR/Western confirmed expected bPML differences, and IFA showed corresponding changes in PML-NBs (Figure 6C–E). Following infection at MOI = 1, quantification of viral gB copies at 0.5–12 hpi showed that PML^Hi-MDBK cells accumulated significantly fewer viral genomes than parental MDBK and shPML-MDBK cells (P < 0.05), whereas shPML-MDBK tended to harbor more viral DNA than MDBK, though not statistically significant (Figure 6F). Thus, IFN-α enhances bPML abundance and NB organization, and elevated bPML constrains very-early BoHV-1 replication.

**Figure 6.**
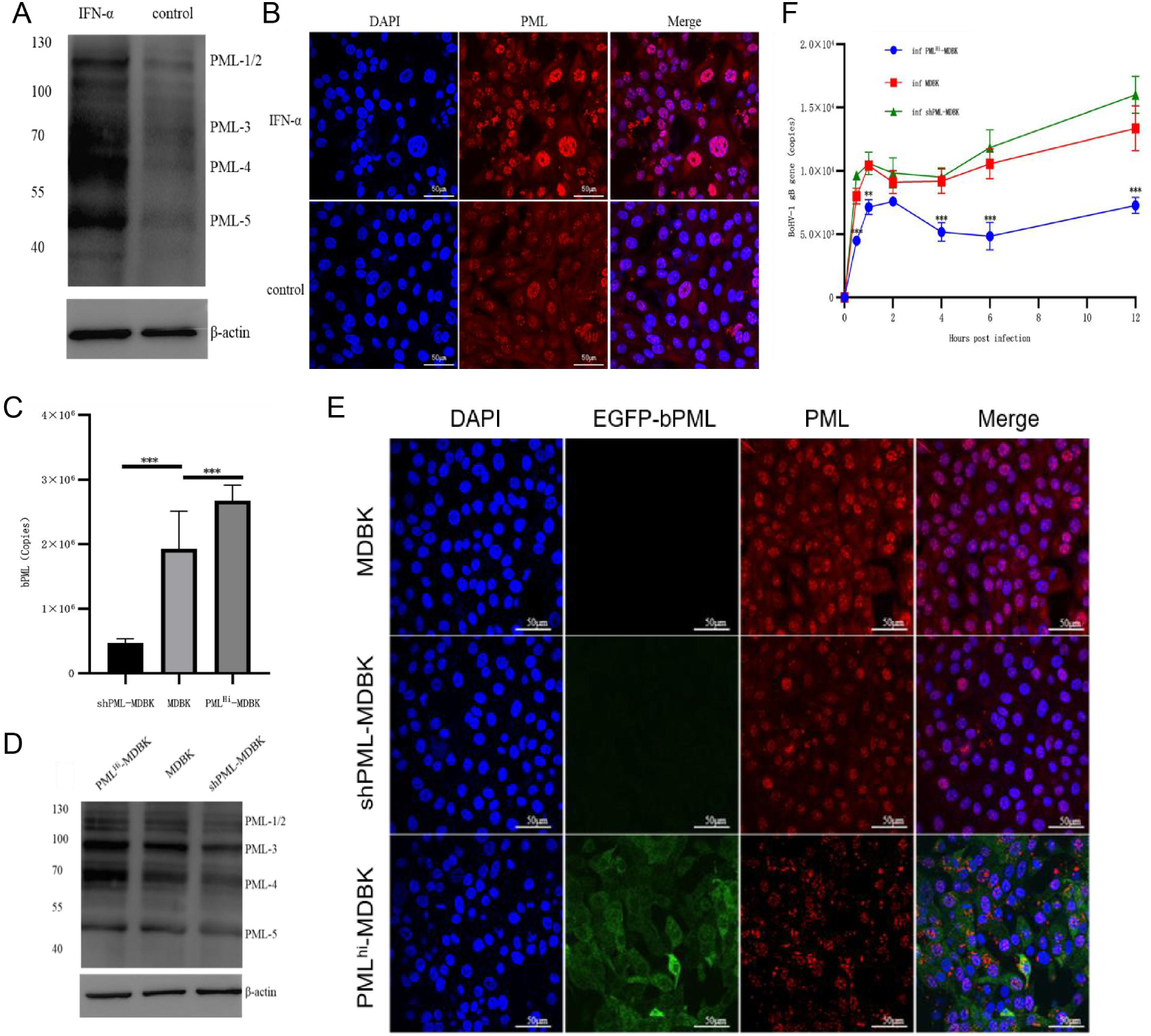
IFN-α alters bPML/PML-NBs and constrains early BoHV-1 replication measured by kinetic qPCR. (A) Western blot of MDBK ± IFN-α to assess bPML protein. (B) Immunofluorescence (PML, red; nuclei, DAPI; scale bars, 50 µm) to visualize PML-NB remodeling after IFN-α. (C) qRT-PCR of bPML in shPML-MDBK, parental MDBK, and PML^Hi-MDBK to verify knockdown/overexpression. (D) Western blot confirmation of bPML levels in the three lines. (E) Immunofluorescence of shPML-MDBK and PML^Hi-MDBK to examine PML-NB abundance and distribution. (F) Early replication assay: cells infected with BoHV-1 (MOI = 1) and viral gB DNA quantified by qPCR at 0.5–12 h post-infection. Data are mean ± SD (n = 3); statistical tests and thresholds are provided in Methods.

### 3.6. BoHV-1 subverts bPML restriction by dismantling PML-NBs

To test whether BoHV-1 attenuates bPML-mediated restriction by targeting PML-NBs, we first generated and validated bPML-specific reagents. Recombinant bPML produced in E. coli (pET-32a-bPML/Transetta(DE3)) migrated at ∼55 kDa by SDS–PAGE; Ni-NTA chromatography yielded a single predominant species, and immunoblotting confirmed identity with anti-His and anti-bPML antibodies (Supplementary Figure 2A–D). Using the purified antigen, we raised rabbit/mouse polyclonals and obtained a monoclonal antibody (bPML-2G5) by hybridoma screening; bPML-2G5 specifically recognized exogenous bPML in 293T cells by IF and immunoblot (Supplementary Figure 3A–C). With these tools, IF of BoHV-1–infected MDBK cells at the onset of CPE showed loss of discrete nuclear PML-NB puncta, fragmentation/dispersion of NB structures, and partial relocalization of bPML to the cytoplasm (Figure 7A). In vivo, double IF of trigeminal ganglion sections demonstrated nuclear co-localization of bPML with ICP0 at 4 dpi, whereas by 14 dpi the NB architecture was disorganized with fewer, irregular puncta and a more diffuse nuclear signal (Figure 7B). Complementary protein–protein docking supported a direct bPML–bICP0 interface (top HADDOCK cluster score −53.0 ± 3.5; Figure 7C). Together, these data indicate a bidirectional antagonism: intact PML-NBs contribute to early restriction of BoHV-1 by bPML, and the virus counters by dismantling PML-NB integrity.

**Figure 7.**
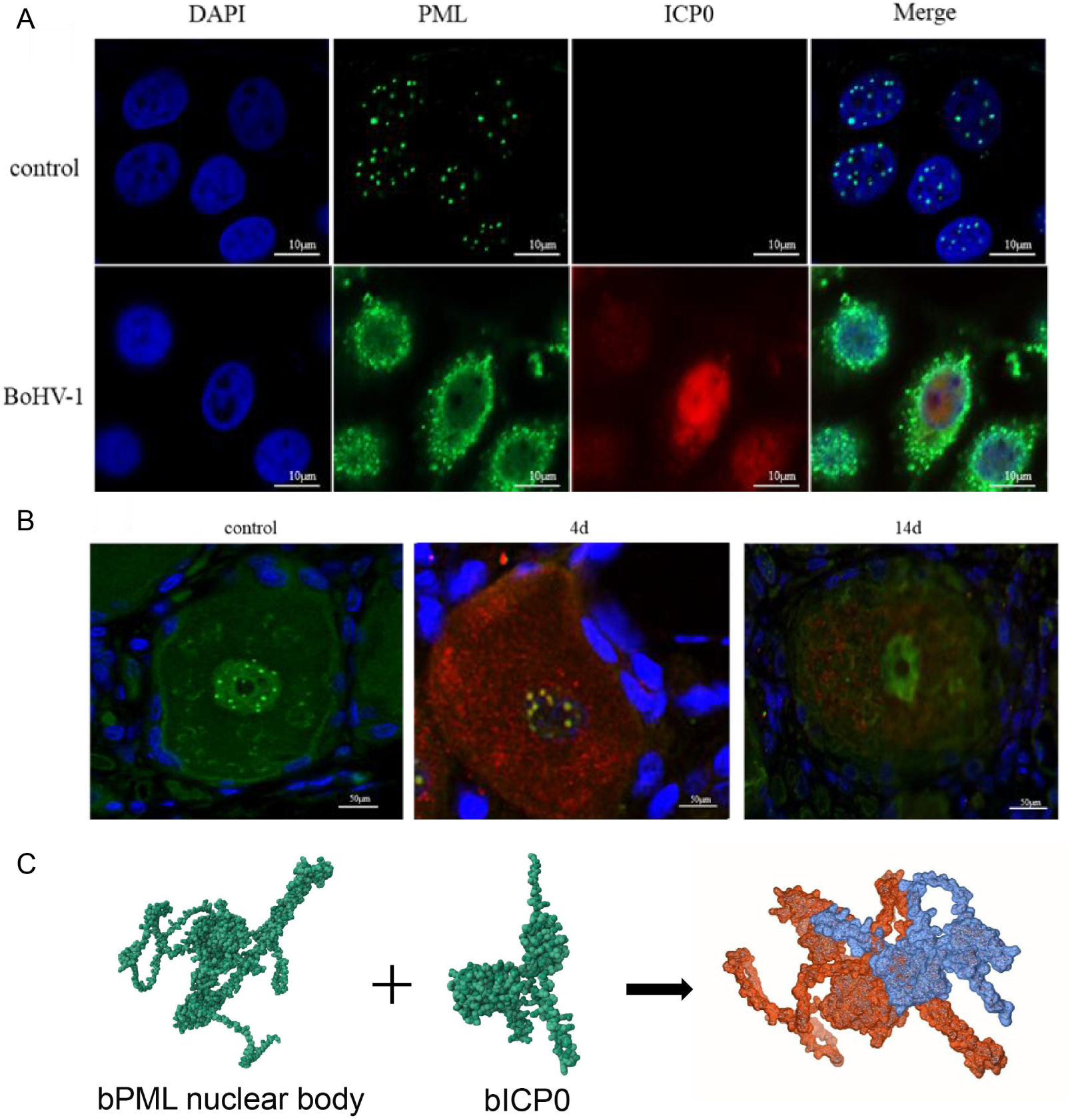
BoHV-1 targets PML nuclear bodies in vitro and in vivo, with a plausible bPML–bICP0 interface supported by docking. (A) MDBK cells infected with BoHV-1 (MOI = 1) were fixed at CPE onset and stained with anti-bPML (green), anti-ICP0 (red), and DAPI (blue); scale bars as indicated. (B) Trigeminal ganglia from infected calves were collected at 4 and 14 dpi, sectioned, and subjected to double IF with anti-bPML (green) and anti-ICP0 (red) plus DAPI counterstain (blue); scale bars as indicated. (C) Protein–protein docking: the bPML–bICP0 interaction was modeled using HADDOCK; the top cluster (n = 30) yielded a score of −53.0 ± 3.5 and a buried surface area of 994 ± 67 Å². Docking parameters and evaluation criteria are detailed in Methods.

### 3.7. Isoform-specific modulation of early BoHV-1 replication by bPML

To determine isoform specificity, Vero cells were transiently transfected with bPML1–bPML6 and infected (MOI = 1). Viral immediate-early (bICP0) and structural (gB) gene copies were quantified at 1, 3, and 6 hpi. Isoforms exhibited clear bidirectional effects: bPML1 and bPML6 consistently increased bICP0/gB copies (strongest at 3–6 hpi; Figure 8A,F,G,L), bPML2 produced a modest increase (Figure 8B,H), whereas bPML3, bPML4, and bPML5 reduced accumulation across the time course, with bPML5 showing the most stable suppression (Figure 8C–E,I–K). Differences were minimal at 1 hpi and became pronounced by 3–6 hpi, indicating isoform-specific, opposing control over very-early genome accumulation.

**Figure 8.**
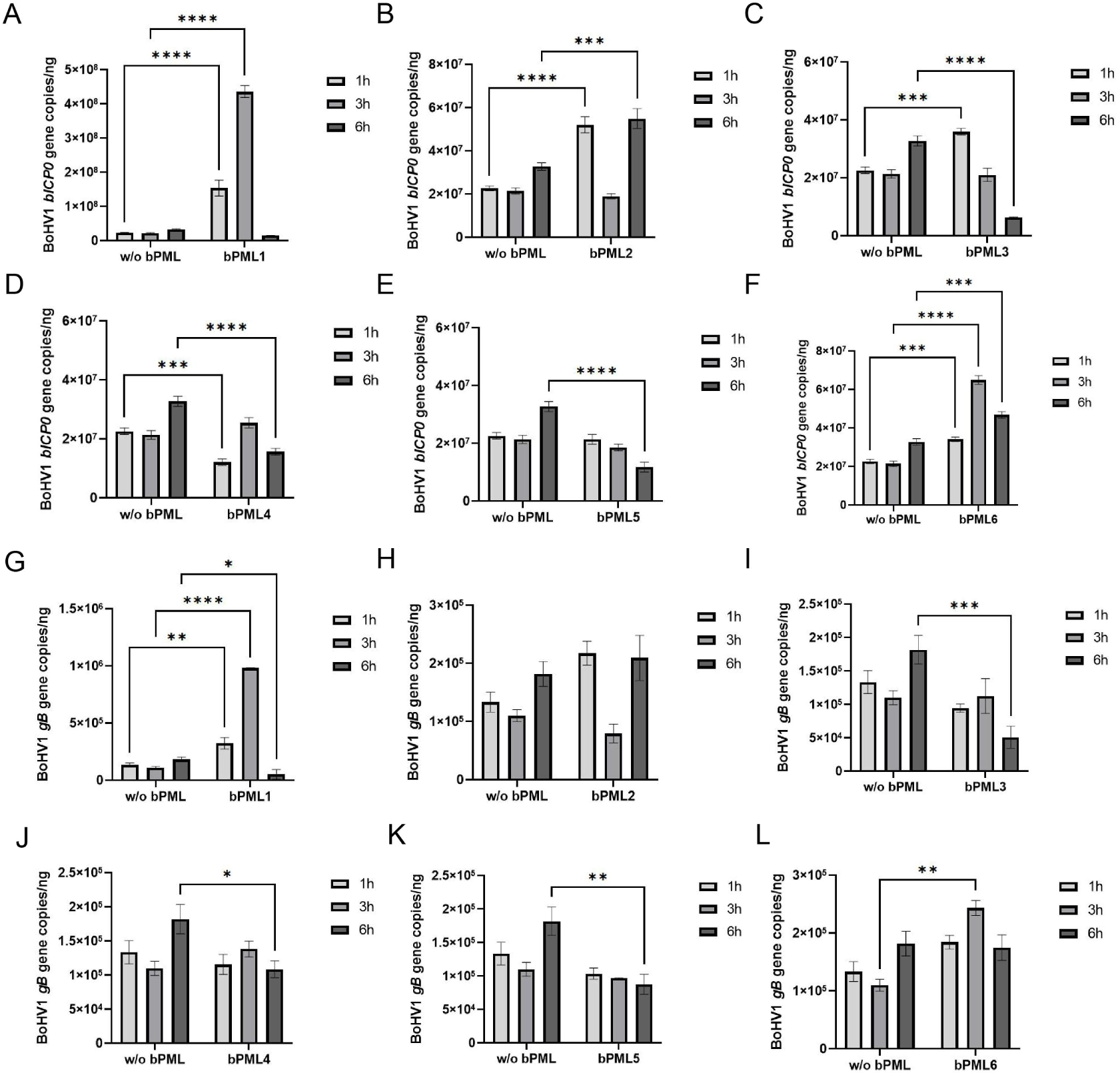
bPML isoforms differentially modulate early BoHV-1 gene accumulation. Vero cells transiently expressing individual bPML isoforms (bPML1–bPML6) or vector control (w/o bPML) were infected with BoHV-1 (MOI = 1). At 1, 3, and 6 hpi, qPCR quantified bICP0 (A–F)and gB (G–L)gene copies (mean ± SD; significance indicated on plots). Statistical tests and replication details are provided in Methods.

## 4. Discussion

Our controlled calf infection delineates a coherent sequence in BoHV-1 pathogenesis. High-titer shedding from upper-airway mucosa (nasal > ocular ≫ rectal) peaked at ∼3–6 dpi and contracted by 10–14 dpi; tissue distribution was dominated by tonsil/URT at 4 dpi with a lower yet detectable TG signal; and TG remained positive at 14 dpi despite falling mucosal loads, indicating neuroinvasion and the onset of persistence [2, 14, 18-20]. Transcriptomically, TG exhibited a two-phase program—limited sensing/proteostasis priming at 4 dpi (COG K/O/T/I) and broad immune–ECM activation at 14 dpi (COG V/O/T with D/L/U/Z)—with GSEA enrichment of chemotaxis and complement/humoral modules and repression of synaptic programs, mirroring tissue virologic kinetics [4, 21, 22].

Host proteostasis emerges as a unifying mechanism for both antiviral restriction and viral counteraction [23, 24]. At 14 dpi, EggNOG/COG highlighted post-translational modification/protein turnover/chaperones, and GSEA revealed coordinated up-regulation of ISG/antigen-processing and proteasome modules. Because PML-NB integrity requires SUMOylation and poly-SUMOylated PML is recognized by SUMO-targeted ubiquitin ligases (e.g., RNF4) for proteasomal degradation, enhanced PTM/turnover networks could simultaneously amplify antiviral proteostasis within NBs and create an avenue for bICP0-driven NB disassembly via SUMO–ubiquitin crosstalk [25-27]. Cross-species STRING mapping was concordant with this model: a conserved PML–SUMO1–UBE2I–DAXX–SP100 core linked to TP53–MDM2 and RNF4 in human and bovine networks, with species-specific extensions (e.g., SPI1 in human; EP300 and a bovine SUMO/ubiquitin-linked node) that may tune chromatin states and innate outputs [28-30].

Mechanistically, the IFN-α–PML axis constrains very-early replication, which BoHV-1 subsequently counteracts [31-33]. IFN-α upregulated bPML and enlarged PML-NBs, and bPML elevation reduced BoHV-1 genome accumulation at 0.5–12 hpi, whereas bPML knockdown tended to increase it, supporting PML as an intrinsic, IFN-sensitized antiviral hub [10, 16]. Conversely, BoHV-1 compromised PML-NB integrity: in MDBK cells at CPE onset we observed fragmentation/dispersion of nuclear PML puncta and partial cytoplasmic relocalization; in TG, nuclear bPML/ICP0 colocalization at 4 dpi gave way to disorganized NB architecture at 14 dpi. Docking supported a direct, energetically favorable bPML–bICP0 interface. This bidirectional antagonism echoes the HSV-1 ICP0 strategy—targeting PML/SP100 and dismantling PML-NBs to relieve intrinsic restriction—and argues that bICP0 executes a functionally analogous countermeasure in bovine cells [10, 34].

Isoform-level analyses suggest PML’s antiviral output is modular and tunable [12, 35]. In Vero cells, bPML1/6 (and to a lesser extent bPML2) enhanced early bICP0/gB accumulation, whereas bPML3/4/5 suppressed it, with bPML5 showing the most stable inhibition. Such isoform dependence, reported for human PML in HSV models [36, 37], likely reflects differences in C-terminal SUMO-interacting motifs, partner recruitment (e.g., DAXX/SP100), and NB ultrastructure. Our data extend this principle to bovine PML and BoHV-1, implying that isoforms bias PML-NB composition toward restriction-competent or restriction-deficient states.

These molecular events have clear implications for the TG niche [38, 39]. By 14 dpi, the microenvironment displayed chemokine-rich, ECM-remodeling signatures alongside suppression of neuronal/synaptic programs—an antiviral glial–neuronal milieu that prioritizes leukocyte recruitment and antigen processing over neurotransmission, potentially constraining viral spread at the cost of transient neuronal function. Given PML’s roles at the intersection of IFN signaling, chromatin regulation, and stress pathways [7, 30, 40], PML-NB state may influence how TG balances immune activation with neuronal homeostasis in the late acute phase.

Limitations include modest group sizes (n = 3), which limit power for subtle effects and preclude stratification by cell type; reliance on Vero/MDBK systems that capture conserved mechanisms but lack TG cellular diversity; and focus on 4 and 14 dpi. Intervening/later intervals (e.g., 21–35 dpi) and bona fide reactivation models are needed to map the continuum into latency/reactivation [2, 41]. Viral DNA copies served as readouts; integrating infectious titers and immediate-early/late transcripts would refine replication staging. Finally, STRING-based PPIs infer functional proximity but do not establish direct physical binding for all edges; targeted biochemical validation is essential.

Future work should test whether bICP0’s RING and SIM-interacting surfaces are necessary/sufficient for NB dismantling; define isoform-specific bPML domains that govern restriction and bICP0 sensitivity; perturb SUMO/ubiquitin enzymes (UBE2I, PIAS, RNF4) while monitoring NB dynamics and BoHV-1 output; deploy ChIP-seq/ATAC-seq and proximity proteomics from TG to connect PML-NB states to chromatin and partner composition; and apply single-cell/spatial transcriptomics to assign TG programs to neurons, satellite glia, and infiltrating leukocytes. Translationally, leveraging IFN-α or small-molecule enhancers of PML-NB integrity— while avoiding proteostasis imbalances that favor NB dismantling—may reduce acute replication and limit seeding of latency, pending in vivo efficacy and safety in cattle.

## Authors’ contributions

J.C., M.C., Y.W. carried out experiments, analysed data, and drafted figures; J.C. and M.C. contributed to the in vivo experiments and data analysis; Y.W. and F.W. performed bioinformatic RNAseq data analyses; Y.L. designed and supervised the study. Y.W. and Y.L. reviewed data, wrote, and revised the manuscript. All authors reviewed, edited, and approved the manuscript.

## Institutional Review Board Statement

All experiments were approved by the Ethical Committee of Beijing Academy of Agriculture and Forestry Sciences (SYXQ-2012-0034).

## Informed Consent Statement

Not applicable.

## Data Availability Statement

The data presented in this study are available on request from the corresponding author. The data are not publicly available due to data are still being processed to produce other papers.

